# The *Porphyromonas gingivalis* lipid A 1-phosphatase LpxE has unique features and requires a functional type IX secretion system for its activity

**DOI:** 10.1101/2024.12.20.629629

**Authors:** Sunjun Wang, Yichao Liu, Beichang Zhang, Joseph Aduse-Opoku, Roberto Buccafusca, Giulia Mastroianni, Pedro Machado, Mark A. J. Roberts, Michael A. Curtis, James A. Garnett

## Abstract

*Porphyromonas gingivalis* is a Gram-negative bacterium that plays a central role in the development of periodontal disease. It uses a type IX secretion system (T9SS) to export a range of virulence factors to the bacterial surface where they are attached to A-LPS, one of the two forms of lipopolysaccharide (LPS) produced in *P. gingivalis*, and then packaged into outer membrane vesicles (OMVs). We previously showed that 1-P dephosphorylation of the lipid A component of LPS is regulated by the T9SS outer membrane protein (OMP) PorV, and this is linked to membrane destabilisation and OMV blebbing/formation. In this study we aimed to extend this and investigate the role of other T9SS OMPs in OMV biogenesis. We examined gingipain activity, gingipain secretion, A-LPS production, OMV morphology, and lipid A structure in *P. gingivalis* W50 and T9SS OMP mutant strains, and our results support an essential role for these proteins in type IX secretion. In addition, we produced a lipid A 1-phosphatase (Δ*lpxE*) mutant and show that all T9SS OMPs are required for LpxE activity and correct vesicle formation. LpxE has a unique C-terminal extension, and we propose that a cargo protein exported by the T9SS can directly/indirectly interact with this and regulate LpxE activity. This study provides insight into a new mechanism that links type IX cargo sorting with OMV blebbing, which may also be present in other Bacteroidota that colonise the gut and oral cavity.

## Introduction

Periodontitis is a biofilm-associated inflammatory disease which affects the supporting tissues around the teeth and is linked with a range of chronic diseases including cardiovascular disease, diabetes, rheumatoid arthritis, and pregnancy complications (1, 2). The global incidence of severe periodontitis is estimated at 11% and it is ranked as the sixth most prevalent disease worldwide. *Porphyromonas gingivalis* is a major microbial agent of periodontitis (3), and it uses a type IX secretion system (T9SS) to export key virulence factors onto the bacterial surface (4, 5). Subsequently these can be sorted into outer membrane vesicles (OMVs) (6), which are spherical structures that bleb from the outer membrane and can transport virulent traits long distances from their source (7). For example, the lysine-specific (Kgp) and arginine-specific (RgpA, RgpB) gingipain cysteine proteases are exported to the *P. gingivalis* surface and OMVs (8), where they carry out a range of functions that include host adhesion, modulation of host immunity, and degradation of host proteins and tissues (9).

The T9SS is produced by a wide range of bacteria across the Fibrobacteres-Chlorobi-Bacteroidetes superphylum and is highly versatile. While in *P. gingivalis* its main function is the export of virulence factors, in *Flavobacterium johnsoniae* it enables gliding motility (10), and in *Tannerella forsythia* it supports S-layer formation (11). The T9SS is composed of two major structures: a >1.4-MDa “translocon complex”, which crosses both the inner and outer membranes (12, 13), and a smaller “attachment complex”, located in the outer membrane (14). Within the translocon complex, PorL forms a pentamer in the inner membrane which can generate proton motive force and rotate a dimer of PorM, which extends across the periplasm (15–17). A dodecamer of PorL/PorM sub-complexes are arranged as a tube-like structure, with PorM binding PorN, which forms an oligomeric ring with PorK on the periplasmic face of the outer membrane (13, 18). Above the PorK/PorN ring on the extracellular surface there are eight copies of the tranlocon pore protein, Sov (13). These associate with a β-barrel outer membrane protein (OMP), PorV, a periplasmic lipoprotein, PorW, and a periplasmic protein, SkpA, which anchor Sov to the PorK/PorN ring, and bind a protein of unknown function, PorD, respectively (13, 19–21).

All T9SS cargo proteins contain a conserved Ig-like C-terminal domain (CTD) (22–25), and after entering the periplasm via the SEC pathway, the CTD is directed into the Sov channel through interactions with PorM, PorN, SkpA, PorW and PorV (20, 26). Although the precise mechanism is unclear, it has been proposed that transfer of proton motive force from the inner membrane to the PorK/N ring may stimulate the lateral release of cargo proteins from Sov (20). This results in the association of a periplasmic plug protein inside Sov to prevent leakage from the periplasm (21) and release of a binary PorV/cargo complex in the outer membrane (25), which acts as a shuttle to move cargo proteins away from Sov to an attachment complex (14). The attachment complex is formed from PorQ, PorU, PorV, and PorZ in a 1:1:1:1 stoichiometry. PorQ is another OMP, and PorQ and PorV tether PorZ and PorU, respectively, to the bacterial surface (12, 14, 27).

Lipopolysaccharide (LPS) is the major component of the outer membrane of Gram-negative bacteria, and it is composed of lipid A, a core oligosaccharide, and a distal O-antigen of repetitive glycan polymers (28). *P. gingivalis* produces two forms of LPS, the conventional neutral O-antigen polysaccharide linked LPS (O-LPS) and anionic-LPS (A-LPS) which has an anionic O-antigen polysaccharide (29). Within the attachment complex, PorZ has a role in presenting A-LPS to PorU, a sortase, which then cleaves off the cargo CTD and mediates A-LPS attachment to the C-terminus of the truncated cargo via a short carbohydrate linker (30–34).

In addition, other OMPs have also been reported to be essential for correct type-IX secretion. PorP is an OMP that interacts with the PorK/PorN ring and through association with the periplasmic lipoprotein PorE, it is thought to anchor the T9SS to peptidoglycan (35, 36). In addition, PorP binds type-B CTDs/cargo and attaches them to the bacterial surface in an A-LPS independent manner (35). Likewise, PorG, PorT and PorF are also OMPs that are essential for correct functioning of the T9SS, but while PorG has been shown to interact with the PorK/PorN ring (18, 37), the interaction profiles of PorT and PorF are unknown (22).

Many bacteria can express lipid A phosphatase and/or *O*-deacylase enzymes in their inner and outer membranes, respectively, which promote bacterial evasion of host immunity but also alters the properties of their outer membrane (28). In *P. gingivalis*, one lipid A 1-phosphatase (LpxE/PG1773), two lipid A 4-phosphatase (LpxF1/PG1587, LpxF2/PG1738), and one 3-*O*-deacylase (LpxR/PG1333) have been identified. These have been implicated in *P. gingivalis* evasion of TLR4 sensing through the production of dephosphorylated/deacylated forms of lipid A (38–41). However, we have also shown that deletion of *porV* in *P. gingivalis* W50 strain results in abrogation of lipid A 1-phosphatase activity, and deformation of OMVs (42). In this study we aimed to understand whether the regulation of LpxE activity was specific to PorV function, or whether this was a more general consequence of type-IX secretion at the outer membrane. We demonstrate that deletion of other type-IX associated OMPs in *P. gingivalis* W50 also cause defects in T9SS function and LpxE activity and production of larger/deformed OMVs. A Δ*lpxE* mutant strain also produced distorted vesicles and displayed minor defects in RgpA and/or RgpB activity, but not Kgp activity. *P. gingivalis* LpxE has a distinct sequence, containing an additional 25 kDa extension at its C-terminus (39), and we have identified similar LpxE enzymes in other *Bacteroides* species that colonise the gut and oral cavity. We propose that in *P. gingivalis* this unique C-terminal region can interact directly/indirectly with the T9SS to regulate LpxE activity, and destabilisation of the outer membrane through lipid A modification/A-LPS repulsion is important for OMV development.

## Materials and Methods

### Bacterial strains and media

All strains used in this study are listed in **Supplementary Table 1**. *P. gingivalis* W50 strain served as wild type and parent for all mutants and were routinely grown on blood agar plates containing 5% (v/v) defibrinated horse blood or in brain heart infusion (BHI) broth supplemented with hemin (5 μg/ml), in an anaerobic atmosphere of 80% N_2_, 10% H_2_, and 10% CO_2_ (Don Whitely Scientific). Clindamycin (5 μg/ml) or tetracycline (1 μg/ml) was added when required. *Escherichia coli* NEB 5-alpha (New England Biolabs) was used for plasmid maintenance and cloning.

### Gene deletions

Single isogenic mutants defective in *porU*, *porQ*, *porZ*, *porP*, *porT*, *porG, porF* and *lpxE* were generated using primer pairs designed to separately amplify the 5′ and 3′ ends of each ORF by PCR (**Supplementary Table 2**). Following purification and digestion with SacI and XbaI, amplicons were ligated to an *erm* cassette, retrieved from plasmid pVA2198 (43) by T4-DNA ligase. The mixture was purified and used as a template in PCR to generate linear chimeric amplicons that comprise *erm* cassette flanked by the 5′ and 3′ regions of the ORF, which were then electroporated into exponential cells of *P. gingivalis* W50 to generate clindamycin resistant mutants by allelic exchange (44, 45). *P. gingivalis* mutant colonies were then screened by PCR to identify erm cassettes that had been inserted in the correct position. For complementation of *lpxE*, an amplicon corresponding to the *lpxE* ORF and an additional 500 bp regulatory upstream sequence was amplified from W50 genome by PCR and cloned into the pUCET1 complementation plasmid (46) using BglII and NotI sites. The plasmid was linearized with XbaI and the flanking *erm* cassette was then used to target the homologous regions in *lpxE* mutant via electroporation and allelic exchange with selection for the tagged *tetQ* on blood agar plates (29).

### Pigmentation assay

*P. gingivalis* W50 and mutant strains were first grown anaerobically for 24 hrs in BHI broth supplemented with hemin (5 μg/ml) at 37°C. These were then streaked onto blood agar plates and incubated anaerobically at 37°C for 7 days.

### Gingipain activity assay

*P. gingivalis* W50 and mutant strains were grown in BHI broth supplemented with hemin (5 μg/ml) in an anaerobic cabinet for 24 hrs. Whole cell cultures were either assayed straight away or were centrifuged at 9,000 g, 4°C for 25 min and the supernatant retained. Arg- and Lys-specific protease activities were then measured at 30°C in a 1.0 ml reaction volume containing 0.1 M Tris-HCl pH 8.0, 10 mM l-cysteine, 10 mM CaCl_2_, with either 0.5 mM Nα-benzoyl-DL-Arg-*p*-nitroanilide (DL-BR*p*NA) or 0.5 mM N-α-acetyl-L-lysine-*p*-nitroanilide (L-AcLys*p*NA) as the substrate, respectively. The reaction was monitored at 405 nm, and enzyme activity was expressed as increase in absorbance/min/OD600nm cell culture at 30°C.

### Bacterial fractionation

*P. gingivalis* W50 and Δ*lpxE* strains were grown in 20 ml of BHI broth supplemented with hemin (5 μg/ml) in an anaerobic cabinet for 48 hrs. Cells were then centrifuged at 17,000 g, 4°C for 15 min and the supernatant (OMV enriched sample) was applied to a 0.2 μm syringe filter to remove any remaining cells.

Cell pellets were then washed twice in 5 ml of 50 mM Tris-HCl pH 8, 50 mM NaCl, 5 mM CaCl_2_ and centrifuged at 900 g, 4°C for 10 min, and after repeating, and then resuspended in 10 ml of PBS with 10 mM EDTA, and heated in a water bath at 60°C for 30 min. After chilling on ice, the cell suspensions were sonicated with intermittent cooling on ice and then centrifuged at 17,000 g at 4°C for 10 min to remove unbroken cells and cell debris. The supernatant was then centrifuged at 48,400 g at 4°C for 60 min and the pellet was resuspended in 50 mM Tris-HCl pH 8, 50 mM NaCl, 5 mM CaCl_2_, 0.5% (w/v) sarcosine, with constant stirring at room temperature for 30 min. After centrifuging at 48,400 g for 30 min, the outer membrane sample was prepared by reconstituting the pellet in 720 μl of 50 mM Tris-HCl pH 8, 50 mM NaCl, 5 mM CaCl_2_, 1% (v/v) Triton X-114.

### Immunoblotting

Samples were run on a NuPAGE 4–12% SDS-PAGE gel (Invitrogen), followed by transfer onto a PVDF membrane using the semi-dry Invitrogen iBlotter and iBlotter transfer blotting solution. The membrane was incubated in 3% (w/v) BSA in TBST at 4°C overnight, washed three times with TBST and then incubated with primary antibody (rabbit Rb7, rabbit mAb CTD or mouse mAb 1B5) (25, 44, 47) diluted 1:1000 in blocking buffer for 2 hours at room temperature. The membrane was then washed three times with TBST for 10 min and incubated with the secondary antibody diluted to 1:2000 at room temperature for 2 hours. After three, 10 min washes with TBST buffer, membranes were treated with enhanced chemiluminescence substrate (ECL; Pierce) before detection by chemiluminescence (BioRad, ChemDoc).

### Nanosight particle analysis

5 ml cultures of *P. gingivalis* W50 and mutant strains were grown in BHI broth overnight, adjusted to OD_600_ 2.0, and then centrifuged at 26,000 g and 4°C for 30 min. The supernatant was filtered with a 0.22-μm filtration apparatus. The filtrate containing OMVs was diluted 10-fold and subjected to Malvern NanoSight LM10 nanoparticle analysis using a laser light source with wavelengths of 405 nm, 532 nm and 638 nm, and particles were tracked and sized.

### Transmission electron microscopy

*P. gingivalis* W50 and derivatives were grown at 37°C in BHI broth supplemented with hemin (5 μg/ml) in an anaerobic cabinet for 24 hrs. These were then fixed in 100 mM phosphate buffer pH 7.0 containing 3% (w/v) glutaraldehyde, 1% (w/v) formaldehyde and 0.5% (w/v) tannic acid, washed with 100 mM phosphate buffer pH 7.0, and then incubated overnight in 100 mM phosphate buffer pH 7.0 containing 2% (w/v) osmium tetroxide. Bacterial cells (10 μl) were then applied to mesh copper grids, prepared with glow discharged carbon support films, incubated for 2 min, washed five times with 50 μl of 1% aqueous uranyl acetate, and then left to dry for 5 min. Electron micrographs were taken using a JEOL 1230 transmission electron microscope operating at 80 kV. Alternatively, cultures were concentrated by centrifugation for 5 min at 2,000 g, loaded in gold plated carriers and then frozen at high-pressure in a Leica EM ICE high-pressure freezer (Leica Microsystems). For ultrastructural analysis, the samples were freeze substituted in a solution of acetone containing 2% (v/v) osmium tetroxide, 0.1% (v/v) uranyl acetate, and 5% (v/v) distilled water in the Leica AFS. The samples remained at −90°C for 10 hrs, and subsequently warmed to −20°C over a period of 12 hrs. Samples were transferred to 4°C for 30 min, followed by washing with anhydrous acetone at room temperature. The samples were infiltrated and embedded in Spurr resin. Sections of 70 nm were collected using a Leica UC7 ultramictrome, followed by post staining with lead citrate. Electron micrographs were then taken using a JEOL JEM 1400 Flash running at 80 kV. Images were then analysed to determine the mean cell and OMV diameters, where OMVs were within 50 nm of blebbing cells.

### Mass spectrometry

Bacteria were cultured for 48 hours in BHI medium containing 5 μg/ml hemin. LPS was isolated using a modified version of the Tri-Reagent protocol for LPS isolation (48). To generate lipid A, dried LPS samples were resuspended in 10 mM sodium acetate pH 4.5 containing 1% (w/v) sodium dodecyl sulphate (SDS). The solution was heated at 100°C for 1 hour, followed by lyophilization overnight. The resulting lipid A pellets were washed once in ice-cold 95% ethanol containing 0.02 N HCl and then three times in 95% ethanol, followed by a final extraction with 1,160 μl of chloroform-methanol-water (1:1:0.9 v/v/v) to remove residual carbohydrate contaminants. The chloroform layer containing the lipid A was dried and used for MALDI-TOF MS. Samples were dissolved in 10 μl of 20 mg/ml 5-chloro-2-mercaptobenzothiazole (CMBT) in chloroform-methanol at 1:1 (v/v), and 0.5 μl of each sample was analysed in both positive- and negative-ion modes on an AutoFlex Analyzer (Bruker Daltonics) (49).

### Bioinformatics analysis

The primary sequence for *P. gingivalis* LpxE (Uniprot ID: B2RLI7) was analysed using the BLASTp server (50) using a threshold of 0.05, Matrix: BLOSUM62, Gap Costs: Existence 11 Extension 1, Compositional adjustments: Conditional compositional score matrix adjustment. Sequences from unique strains with an accession length >450 residues were then compared using the M-Coffee function within the T-Coffee multiple sequence alignment server (51).

## Results

### T9SS OMP mutants are defective in gingipain secretion

We previously constructed a Δ*porV* mutant in *P. gingivalis* W50, and we initiated this study by creating additional knockout mutants in the other known T9SS β-barrel OMPs (i.e., Δ*porQ*, Δ*porP*, Δ*porT*, Δ*porG*, Δ*porF*) (27), but also Δ*porU* and Δ*porZ*, which are integral components of the outer membrane attachment complex (22). To confirm that our mutant strains were defective in type-IX secretion we examined Arg- and Lys-gingipain activities in both whole cell cultures and isolated culture supernatants, the latter containing soluble gingipains and those enriched within OMVs (**Fig. 1a,b**). For both Arg- and Lys-gingipain activities measured from wild-type W50, ∼80% total activity was observed on the bacterial surface, while ∼20% was observed in the supernatant. Conversely, no activity could be detected for the Δ*porV*, Δ*porP*, Δ*porT* and Δ*porG* strains, and there was a ∼90% drop in the whole cell culture and supernatant activities for the Δ*porQ,* Δ*porZ* and Δ*porF* strains compared to W50. Although the Δ*porU* strain displayed whole cell culture activities comparable with wild type W50, this was primarily localised in the supernatant and indicated that this strain was impaired in retaining gingipains on the bacterial surface.

**Figure 1.**
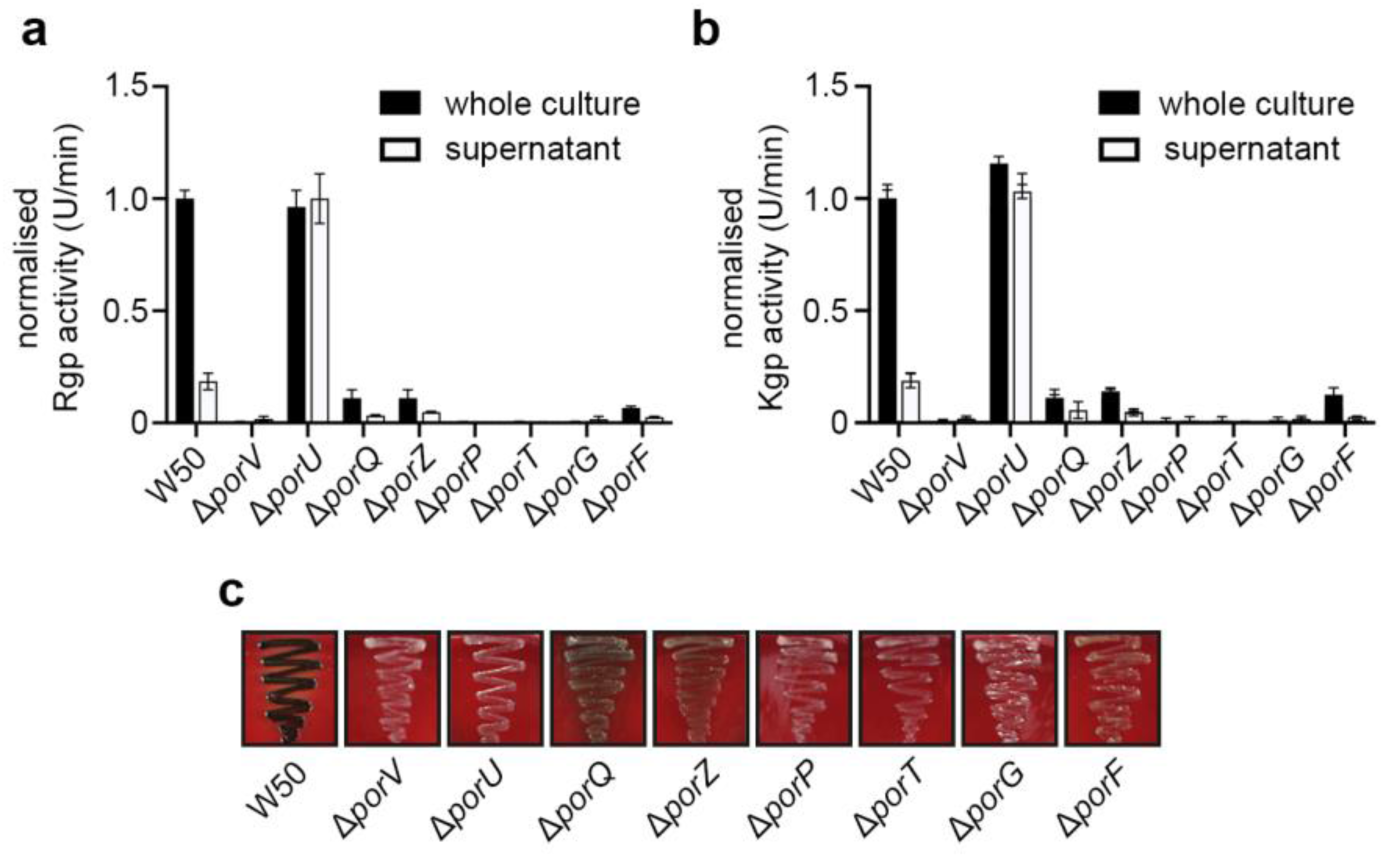
Gingipain activities of *P. gingivalis* T9SS OMP mutants. (a) R-gingipain and (b) K-gingipain activities in whole cultures and supernatants of *P. gingivalis* W50 and isogenic mutant strains, grown in brain-heart infusion broth for 24 hrs. Rgp and Kgp activities were measured using substrates DL-BR*p*NA and L-AcK*p*NA, respectively, and are normalised to WT whole cultures. Data are presented as mean values +/-Standard Error of Mean (SEM) derived from *n* = 2 biologically independent experiments. (c) Pigmentation of *P. gingivalis* strains grown on blood agar plates for 7 days.

We next monitored gingipain export using a colony pigmentation assay. *P. gingivalis* displays black colony pigmentation when grown on blood agar due to Kgp gingipain activity and heme accumulation on the bacterial surface, and when our mutant strains were grown on blood agar, they all exhibited pigmentation defects (**Fig. 1c**). While the Δ*porV*, Δ*porU*, Δ*porP*, Δ*porT* and Δ*porG* strains produced white colonies after 7 days growth, indicative of an inactive T9SS, the Δ*porQ*, Δ*porZ* and Δ*porF* strains instead formed beige colonies, reflecting an impaired export of Kgp with reduced secretion and/or surface attachment.

Strains were then separated into pelleted cells and culture supernatants and immunoblot analysis was then carried out using antisera Rb7, which recognises the RgpA/B catalytic domain (44). In both blots of WT W50, mature RgpB appeared as a diffuse band between ∼70-90 kDa due to its attachment to A-LPS on the outer membrane and OMV surfaces (**Fig. 2a**). We detected no mature gingipains in any of the mutant strains, but instead observed higher molecular weight species which likely correspond to partially processed pro-proteins resulting from defective secretion. Moreover, in the whole cell samples, the Δ*porV*, Δ*porP*, Δ*porT* and Δ*porG* strains displayed similar banding, as did the Δ*porQ* and Δ*porZ* strains, while the Δ*porU* and Δ*porF* strain produced more unique profiles. In the supernatant, the Δ*porV* strain again displayed defective gingipain processing, although RgpB was observed in the Δ*porU*, Δ*porQ* and Δ*porZ* mutants, albeit not attached to A-LPS. No RgpA/B was detected in the supernatant from the Δ*porP*, Δ*porT* and Δ*porG* mutants, while mature RgpB was present in the Δ*porF* strain but at lower levels.

**Figure 2.**
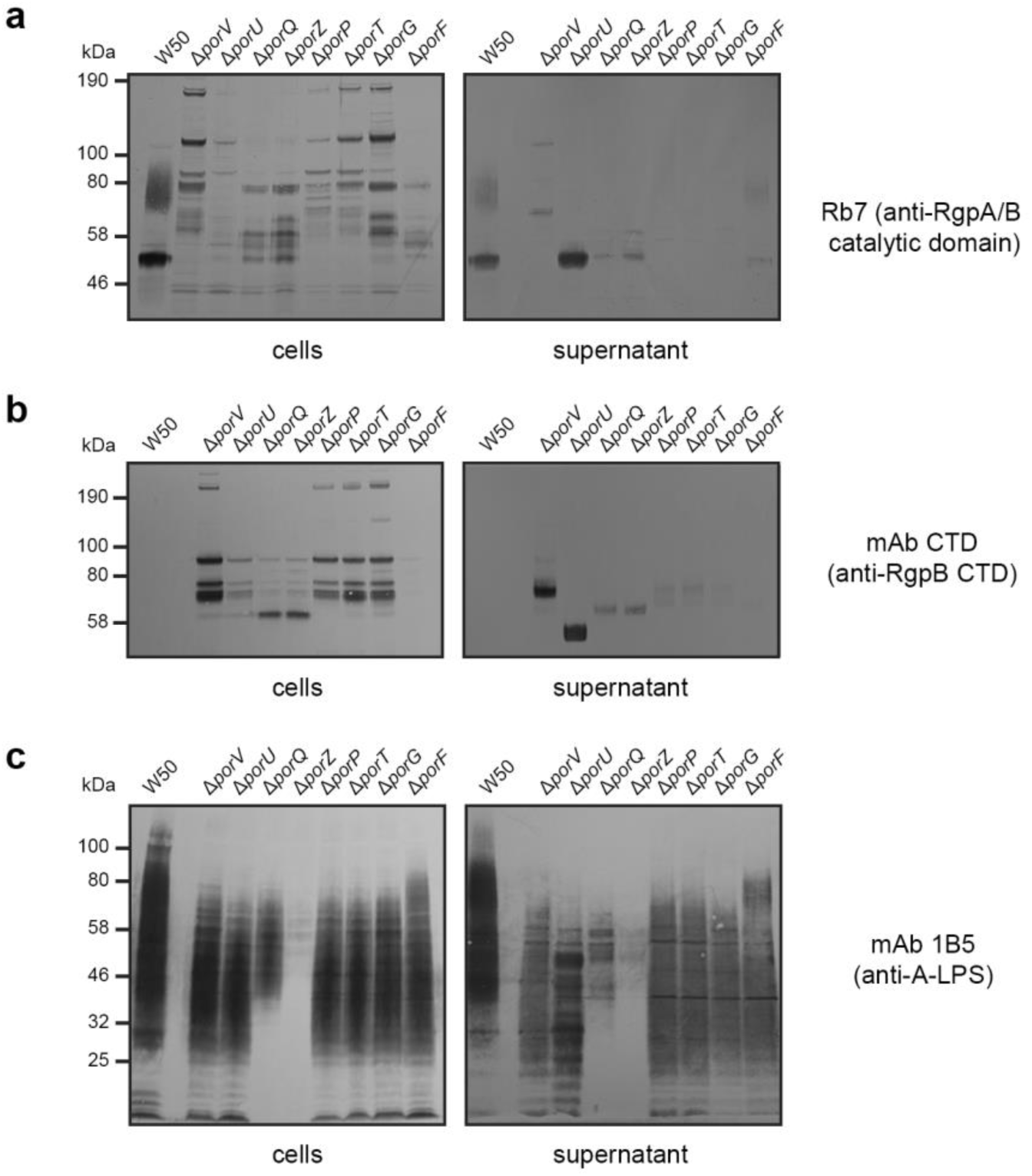
Immunoblots of *P. gingivalis* T9SS OMP mutants. Western blotting of *P. gingivalis* W50 and mutant strain cells and supernatants. (a) Antisera Rb7 against the RgpA/B catalytic domain, (b) mAb CTD against the RgpB CTD and (c) mAb 1B5 against A-LPS.

Immunoblotting was next carried out using monoclonal antibodies against the RgpB CTD (mAb CTD) and the anionic O-antigen of A-LPS (mAb 1B5) (25, 47). Analysis of W50 showed no detection of the CTD in either whole cells or supernatants; likely due to CTD proteolysis upon its release from cargo proteins during A-LPS attachment (**Fig. 2b**). However, in all mutant strains except Δ*porF*, bands were detected at a higher molecular weight to the free CTD (∼9 kDa), and represented partially processed pro-RgpB species, which lack A-LPS attachment and removal of the CTD. In the whole cell blot, while Δ*porU* appeared like W50, Δ*porV*, Δ*porP*, Δ*porT* and Δ*porG* strains, and Δ*porQ* and Δ*porZ* strains, each produced similar band patterns, indicating that proforms of cargo proteins are retained in the cell in these mutants. In the supernatant, only the Δ*porQ* and Δ*porZ* strains, and the Δ*porP*, Δ*porT* and Δ*porG* strains could be grouped together. Moreover, the detection of weak bands in the Δ*porQ*, Δ*porZ*, Δ*porP*, Δ*porT* and Δ*porG* mutants, and strong bands in the Δ*porV* and Δ*porU* mutants indicates that proforms of cargo proteins are released from the cell surface prior to A-LPS attachment or removal of the CTD. Of note, the single band detected in the Δ*porU* mutant supernatant is the expected molecular mass for mature RgpB prior A-LPS attachment. Finally, in the 1B5 blots of W50 cells and supernatants, A-LPS presented as a diffuse band ranging from ∼25-150 kDa (**Fig. 2c**), while in all mutants A-LPS was not detected above ∼75 kDa (potentially A-LPS attached to cargo). However, in both cell and supernatant samples there was an additional decrease in A-LPS species below ∼40 kDa in the Δ*porQ* strain, with an almost complete loss of A-LPS detection in the Δ*porZ* strain. In addition, there was a general decrease in A-LPS detected in the supernatant of all mutants. Together this supports previous observations that the T9SS has a fundamental role in A-LPS transport to the outer membrane (30, 33).

### *P. gingivalis* produce two distinct distributions of OMV sizes

To understand whether these derivative strains also affected the morphology of OMVs as originally observed in the Δ*porV* mutant, we examined them by negative stain transmission electron microscopy (TEM). In line with previous observations (42), a clear electron dense surface layer (EDSL) could be seen on the outer membrane of *P. gingivalis* W50, composed of surface anchored gingipains and capsule, along with OMVs ∼32 nm in diameter blebbing from the outer membrane (**Fig. 3**, **Table 1**). On the other hand, while the average cell diameter was consistent with W50, the EDSL was absent in all mutant strains and their OMVs appeared in general much larger with diameters ranging from ∼99 to 121 nm (**Fig. 3**, **Table 1**). NanoSight single nanoparticle tracking has been used extensively to study vesicle formation in both eukaryotic and prokaryotic systems (52, 53) and we used it here to analyse the size distribution of OMV preparations isolated from overnight culture supernatants of W50, but also strains W83 and ATCC 33277 (**Fig. 4a**). In all three strains we observed a clear bimodal distribution of OMV diameters, with an abundant well-defined species <70 nm in diameter, representing smaller OMVs (modes of 44 nm, 22 nm, and 35 nm, respectively), and a species of larger OMVs with a broader range of diameters from ∼70-200 nm and at lower concentration (modes of 114 nm, 154 nm, and 108 nm, respectively). However, analysis of the W50 mutant strains showed a major loss of the smaller species but with a similar distribution and concentration of the larger OMVs (**Fig. 4b**).

**Figure 3.**
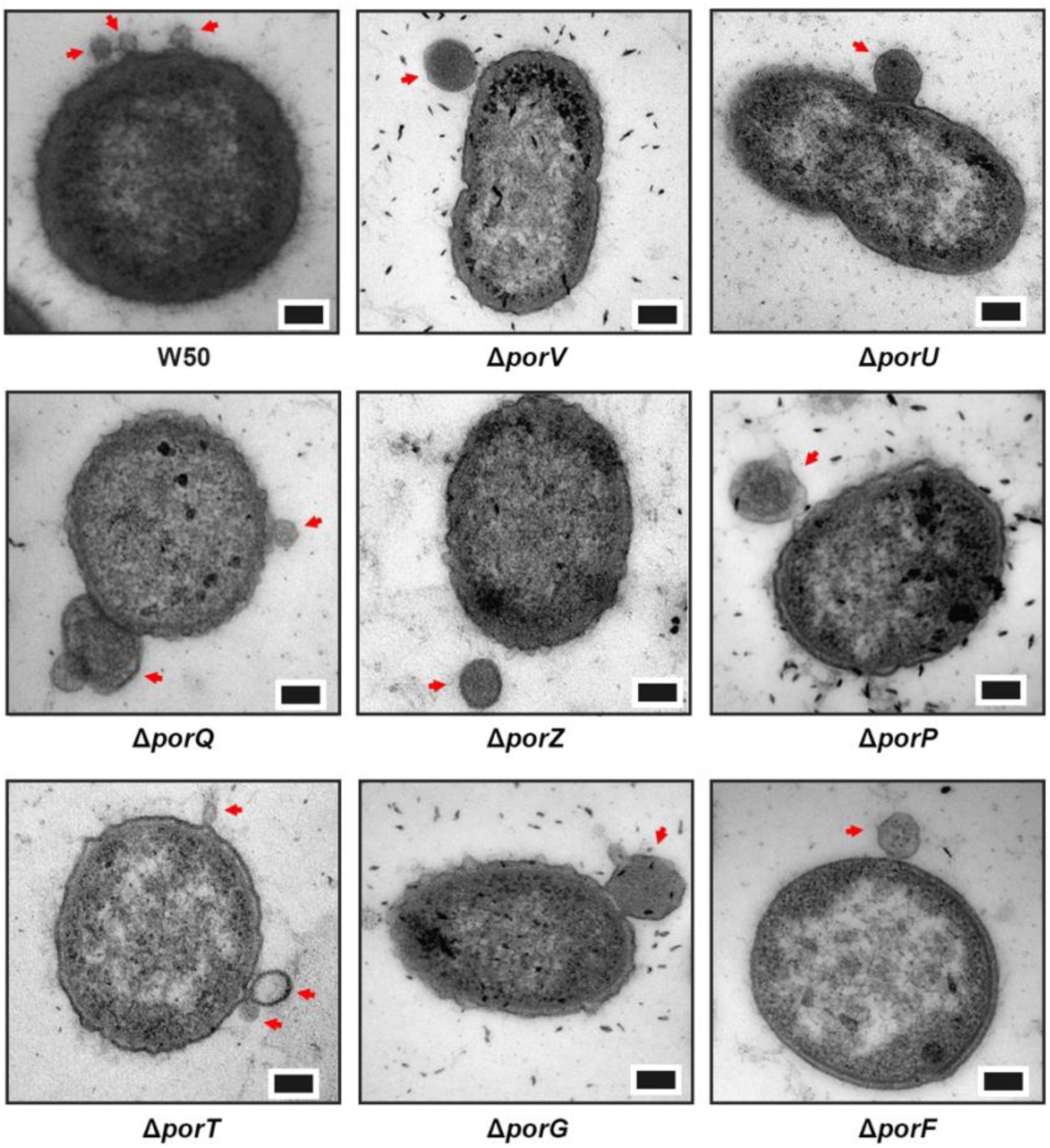
Negative stain TEM of *P. gingivalis* T9SS OMP mutants. Samples were fixed using glutaraldehyde as described in Materials and Methods. The scale bar represents 100 nm. OMVs blebbing from cells are highlighted with a red arrow.

**Figure 4.**
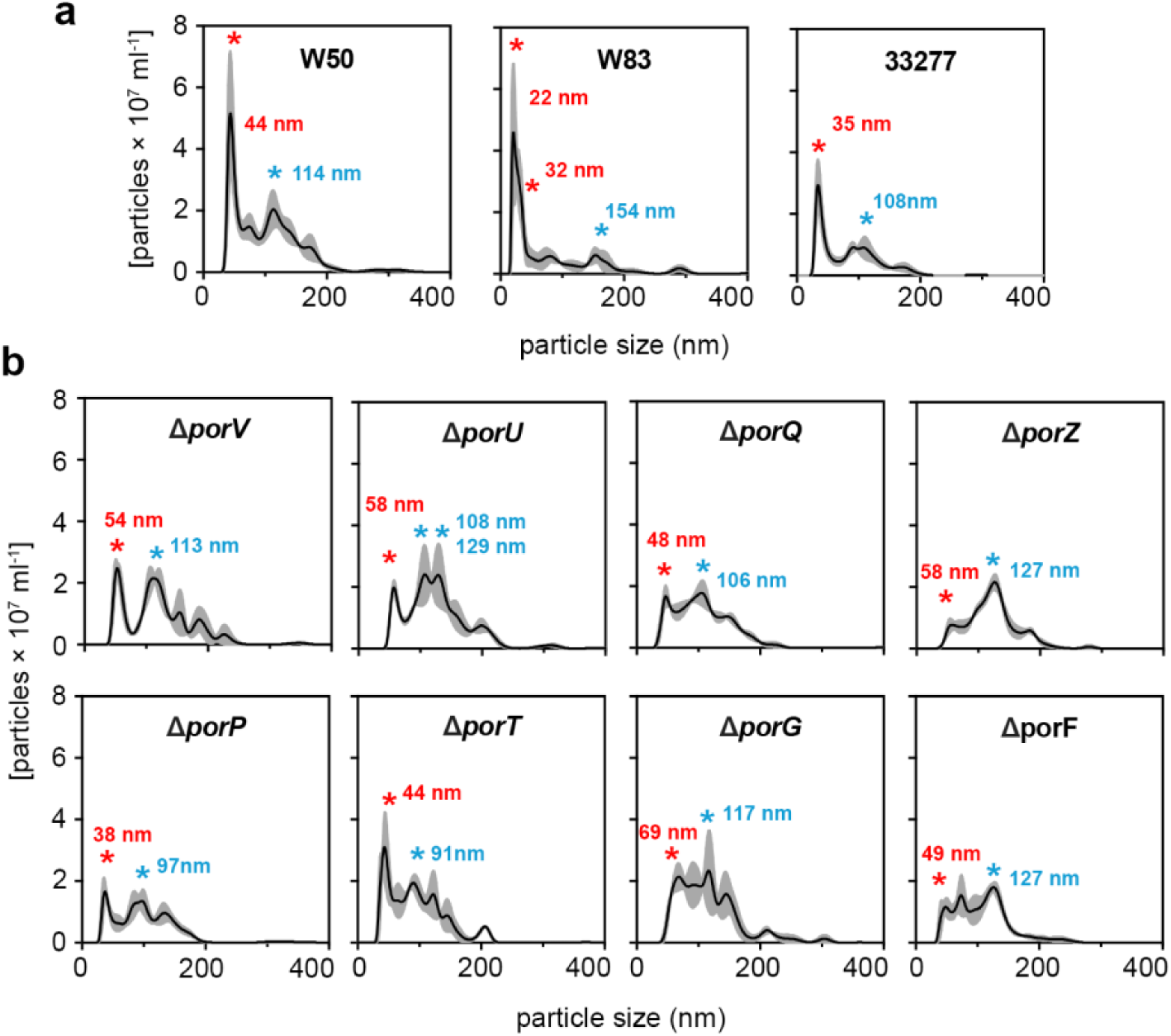
Nanosight analysis of OMVs from *P. gingivalis* W50 and T9SS OMP mutants. NanoSight analysis of crude OMV preparations from (a) W50, W83 and ATCC 33277, and (b) W50 derivative strains. Modes for the smaller and larger OMV species are indicated with a red and blue asterisk, respectively. Data are presented as mean values (black line) +/- Standard Error of Mean (SEM; grey area) derived from *n* = 3 biologically independent experiments.

**Table 1:** Cell and OMV diameters derived from negative stain TEM of formaldehyde fixed *P. gingivalis* W50 and *T9SS OMP* mutants. Data presented for WT and derivative strains as mean values ± standard error of mean (SEM) derived from *n* = 9 bacteria or OMVs.

| | W50 | $\Delta porV$ | $\Delta porU$ | $\Delta porQ$ | $\Delta porZ$ | $\Delta porP$ | $\Delta porT$ | $\Delta porG$ | $\Delta porF$ |
| --- | --- | --- | --- | --- | --- | --- | --- | --- | --- |
| Mean cell diameter (nm) | 449<br>$\pm 30$ | 426<br>$\pm 33$ | 465<br>$\pm 18$ | 434<br>$\pm 27$ | 479<br>$\pm 30$ | 463<br>$\pm 34$ | 442<br>$\pm 32$ | 463<br>$\pm 25$ | 456<br>$\pm 35$ |
| Mean OMV diameter (nm) | 32<br>$\pm 15$ | 111<br>$\pm 20$ | 121<br>$\pm 20$ | 118<br>$\pm 35$ | 109<br>$\pm 16$ | 120<br>$\pm 17$ | 99<br>$\pm 13$ | 116<br>$\pm 15$ | 102<br>$\pm 13$ |

### T9SS OMP mutants display a loss of lipid A 1-phosphatase activity

*P. gingivalis* lipid A is synthesised as a bis-P-penta-acyl form but can be subsequently dephosphorylated and deacylated into mono-P-penta-acyl, mono-P-tetra-acyl, non-P-penta-acyl, and non-P-tetra-acyl species (42). This is due to the activities of LpxF1/2 and LpxE in the inner membrane, which are anticipated to sequentially dephosphorylate bis-P-penta-acyl and mono-1-P-penta-acyl lipid A species, respectively, with subsequent deacylation in the outer membrane by LpxR to generate tetra-acylated mono-1-P and non-P lipid A species (39) (**Fig. 5a**). However, in W50 Δ*porV*, non-phosphorylated species are not detected, which indicates that PorV has a role in regulating the activity of LpxE (39) (**Fig. 5b**). Using MALDI-TOF MS we examined lipid A purified from our new mutant strains and compared them to W50. As before, we detected bis-P-penta-acyl, mono-P-penta-acyl, mono-P-tetra-acyl, non-P-penta-acyl, and non-P-tetra-acyl lipid A species in WT W50, in a ratio of 7:1:9:1:2 while in our Δ*porV* mutant this ratio was 2:3:15:0:0, reflecting a decrease in the bis-P species, and an increase in both mono-P species, but with no detection of the non-P species (**Fig. 5c**). Likewise, analysis of the other T9SS mutants showed a similar phenotype to Δ*porV*, although in the Δ*porQ* and Δ*porG* strains there was also no detection of the bis-P form.

**Figure 5.**
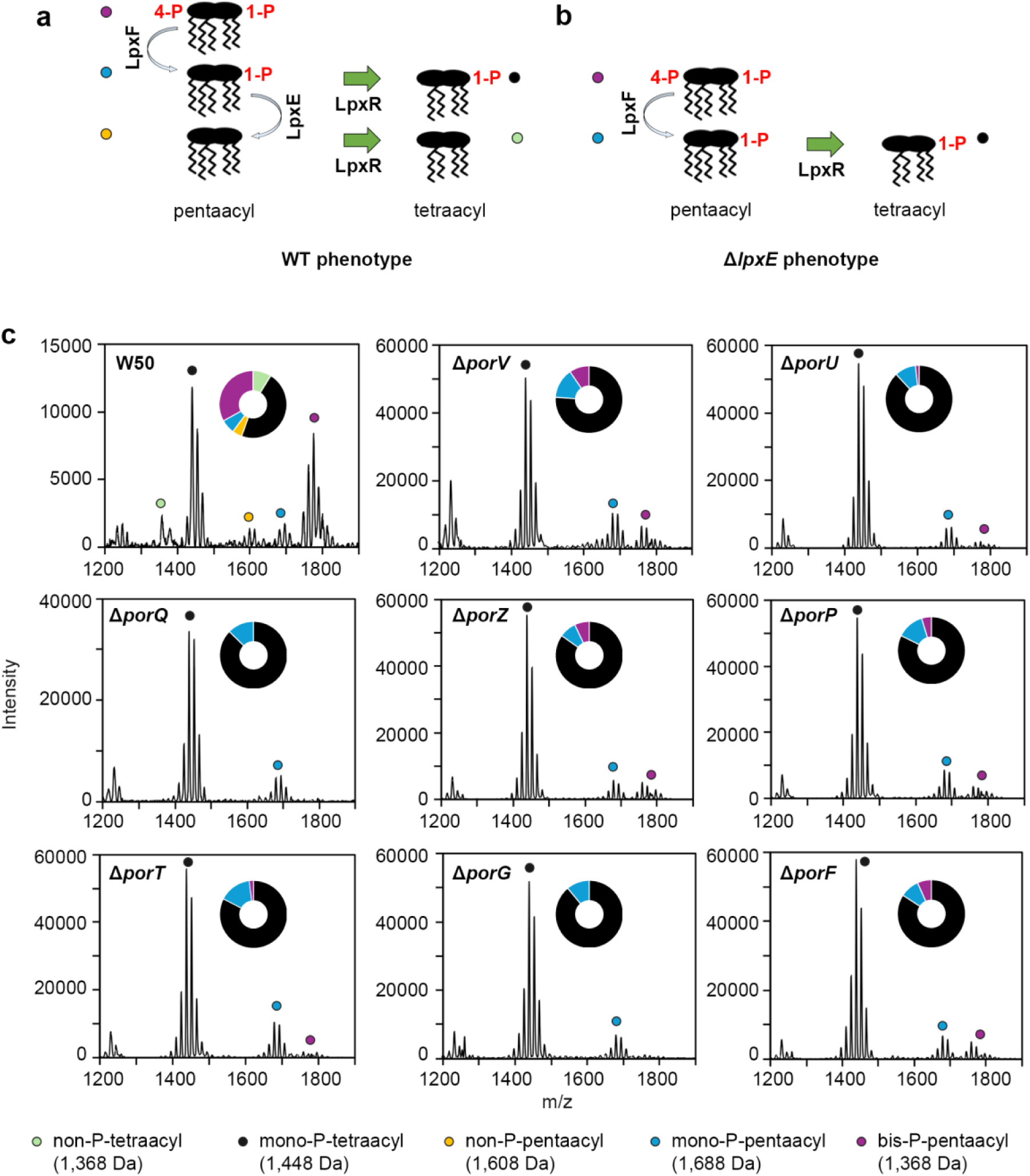
Lipid A modification in *P. gingivalis* T9SS OMP mutants. (a) Schematic showing the lipid A dephosphorylation and deacylation pathway in *P. gingivalis* and (b) Δ*lpxE* mutant. (c) Negative-ion MALDI-TOF MS of lipid A samples highlighting the bis-P-pentaacyl, mono-P-pentaacyl, mono-P-tetraacyl, non-P-pentaacyl, and non-P-tetraacyl species. Wheel chart shows relative amount of each species.

### LpxE affects vesicle formation and cargo sorting

We next created a Δ*lpxE* knock-out in W50 and repeated the above phenotyping. Unlike the T9SS OMP mutants, the Δ*lpxE* strain showed no defect in black colony pigmentation (**Fig. 6a**), in RgpB secretion (**Fig. 6b**), A-LPS biogenesis (**Fig. 6c**) or Lys-gingipain activity in whole cell cultures or isolated culture supernatants (**Fig. 6d**). However, we did observe a ∼35% and ∼25% reduction in the Arg-gingipain activity in whole cell cultures and isolated culture supernatants, respectively, with activity partially recovered upon complementation of the Δ*lpxE* strain with *lpxE* (**Fig. 6e**). We then assessed whether deletion of *lpxE* affected protein accumulation in both the outer membrane and within isolated OMVs. Analysis by SDS-PAGE showed some clear differences and indicated that the function of LpxE also affects the accumulation of specific proteins in both *P. gingivalis* outer membranes and OMVs (**Fig. 6f**). Analysis of OMVs isolated from the Δ*lpxE* strain using NanoSight also showed a major loss of the smaller species but not the larger OMVs (**Fig. 6g**). During the previous negative stain TEM analysis of W50 and T9SS OMP mutants, samples had been fixed using aldehyde cross-linking, which could exacerbate OMV deformation. Therefore, when repeating our TEM analysis with the Δ*lpxE* mutant, we prepared samples using high-pressure freezing and used Δ*porV* as a control (**Fig. 6h**). As before the average cell diameters were consistent with W50, with blebbing OMVs from W50 and Δ*porV* displaying slightly larger diameters of ∼40 nm and ∼120 nm, respectively, however, OMVs blebbing from the Δ*lpxE* strain were much larger with a diameter of ∼164 nm (**Fig. 6h**, **Table 2**).

**Figure 6.**
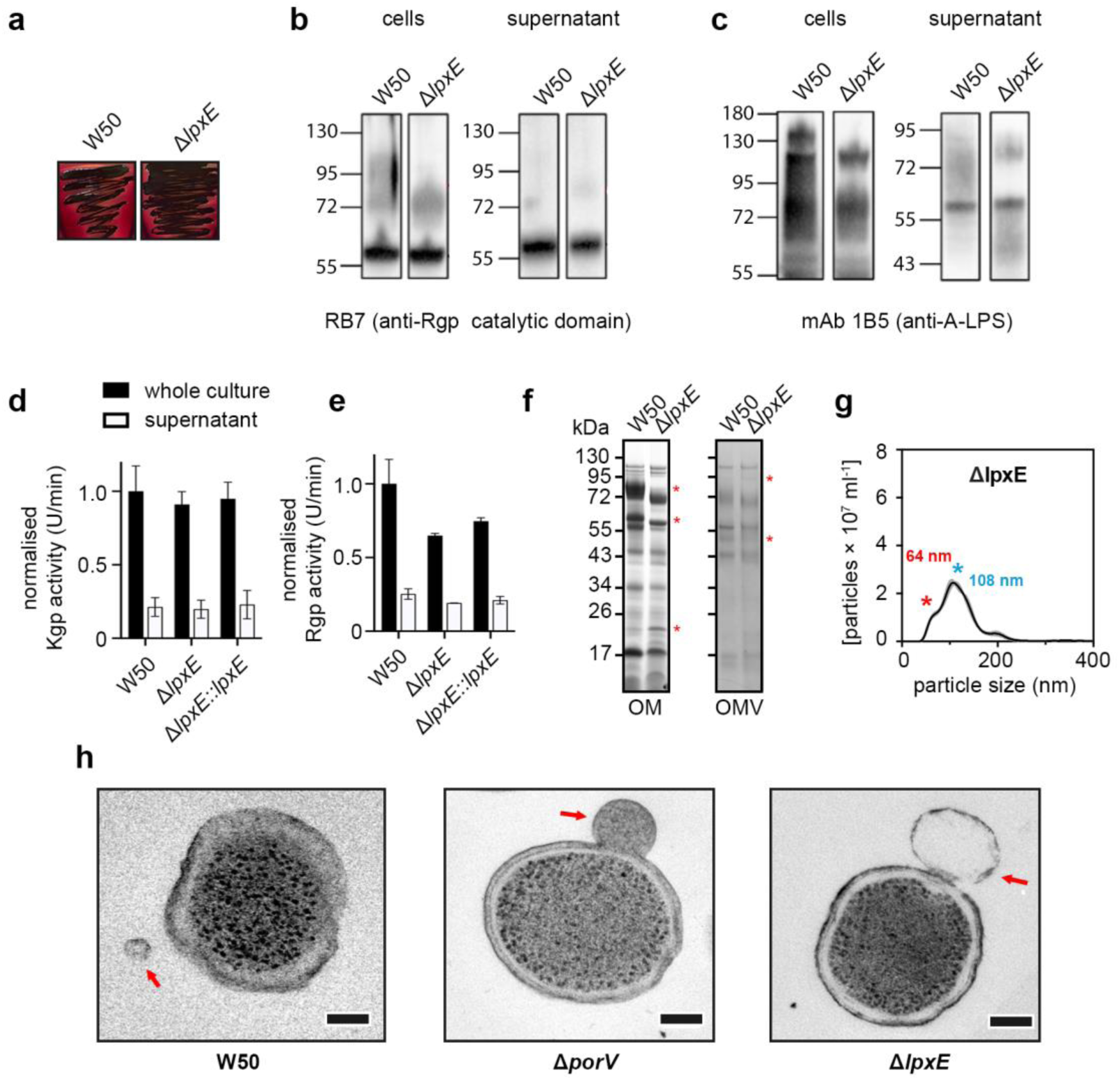
Characterisation of *P. gingivalis* W50 *lpxE* mutant. (a) Pigmentation of WT W50 and Δ*lpxE* mutant grown on blood agar plates for 7 days. (b) Western blotting of *P. gingivalis* W50 and *lpxE* mutant cells and supernatants using antisera Rb7 against the RgpA/B catalytic domain or (c) mAb 1B5 against A-LPS, after growth in brain-heart infusion broth for 24 hrs. (d) K-gingipain and (e) R-gingipain activities in whole cultures and supernatants of *P. gingivalis* W50 and *lpxE* mutant, grown in brain-heart infusion broth for 24 hrs. Rgp and Kgp activities were measured using substrates dl-BR *p* NA and L-AcK*p* NA, respectively, and are normalised to WT whole cultures. Data are presented as mean values +/- Standard Error of Mean (SEM) derived from *n* = 2 biologically independent experiments. (f) SDS-PAGE of outer membranes and OMVs isolated from W50 WT and Δ*lpxE* mutant. Differences between each strain are highlighted with red asterisk. (g) NanoSight analysis of crude OMV preparation from W50 and Δ*lpxE* strains. Modes for the smaller and larger OMV species are indicated with a red and blue asterisk, respectively. Data are presented as mean values (black line) +/- Standard Error of Mean (SEM; grey area) derived from *n* = 3 biologically independent experiments. (h) Negative stain TEM of W50 WT, Δ*porV* and Δ*lpxE* using high-pressure freezing as described in Materials and Methods. The scale bar represents 100 nm. OMVs blebbing from cells are highlighted with a red arrow.

**Table 2:** Cell and OMV diameters derived from negative stain TEM of high-pressure frozen *P. gingivalis* W50 and *T9SS OMP* mutants. Data presented for WT and derivative strains as mean values ± standard error of mean (SEM) derived from *n* = 9 bacteria or OMVs.

| | W50 | $\Delta porV$ | $\Delta lpxE$ |
| --- | --- | --- | --- |
| Mean cell diameter (nm) | 434<br>$\pm 32$ | 448<br>$\pm 20$ | 460<br>$\pm 24$ |
| Mean OMV diameter (nm) | 40<br>$\pm 23$ | 120<br>$\pm 12$ | 164<br>$\pm 14$ |

## Discussion

Type IX-dependent secretion is a highly complex process and involves at least three stages: (i) cargo recognition in the periplasm and passage through the translocon complex, (ii) cargo release and movement across the outer membrane, and (iii) association with the attachment complex and linkage to A-LPS (22). While the role of PorV, PorU, PorQ and PorZ have been relatively well characterised in relation to the second and third stages, the functions of PorP, PorT, PorG and PorF are unclear (22). Previous observations in *P. gingivalis* strains W83 and 33277 have demonstrated that PorV, PorU, PorZ, PorT and PorF are essential for gingipain secretion, with deletion of these genes resulting in white colonies forming on blood agar (30, 33, 54–58). However, here we have shown that deletion of *porQ*, *porZ* or *porF* in W50 instead results in beige coloured colonies forming on blood agar, although Δ*porV,* Δ*porU*, Δ*porP*, Δ*porT* or Δ*porG* mutants form white colonies, suggesting there may be some variation in the absolute level of decrease in protease secretion/maturation in the former strains.

Our examination of Arg- and Lys-gingipain activities in these mutants also showed a similar phenotype, with no activity detected in the Δ*porV*, Δ*porP*, Δ*porT* and Δ*porG* W50 strains, and high loss of activity in Δ*porQ*, Δ*porZ* and Δ*porF* strains, similar to previous reports forΔ *porT* and Δ*porF* mutants in W83 (58, 59). However, our Δ*porU* mutant displayed full gingipain activity in the supernatant but not in whole cells, suggesting that it is able to export gingipains, but these are not retained on the cell surface consistent with its role as a sortase required for A-LPS attachment. A previously reported Δ*porU* mutant in 33277 strain had no gingipain activity, indicating complete loss of secretion (54). This is likely due to differences in how the insertion cassettes are designed, resulting in different partial genes being expressed in each of these studies, although both mutants still support the essential role of PorU. We also carried out immunoblot analysis and again our T9SS OMP mutants displayed major secretion defects with an accumulation of pro-gingipains in the cell and supernatant, similar to that described previously for Δ*porV*, Δ*porU* and Δ*porT* mutants in the 33277 strain (54, 55).

Disruption of *porV* in W50, *porV* or *porT* in 33277 and *porV*, *porZ* or *porZ* in W83 have all been shown to affect A-LPS biogenesis, with an overall reduction in high-molecular weight A-LPS species (A-LPS attached to cargo), and accumulation of lower-molecular weight species in the periplasm (33, 55).

Furthermore, higher levels of A-LPS periplasmic retention were observed in the W83 *porZ* mutant compared with the Δ*porV* and Δ*porT* mutant strains, but with no A-LPS detected in the supernatant containing OMVs (33). Here we show that disruption of T9SS OMPs in W50 presents a similar profile, with a loss in high-molecular weight A-LPS species, an overall reduction of A-LPS in the Δ*porQ* and Δ*porZ* mutants, and an overall reduction of A-LPS in the supernatant. This suggests that PorZ is directly coupled to A-LPS transport to the outer membrane and implies that the primary function of A-LPS is to anchor type IX cargo to the bacterial surface.

Taken together, these data indicate that some of the T9SS OMPs have roles during similar stages of secretion/attachment (**Table 3**). As expected, the Δ*porV*, Δ*porU*, and Δ*porQ/Z* mutants presented distinct phenotypes, reflecting functions related to cargo shuttling and sortase targeting, sortase function, and A-LPS targeting, respectively. On the other hand, the Δ*porP*, Δ*porT* and Δ*porG* mutants have similar phenotypes, and are more likely associated with upstream events. The Δ*porF* mutant was unique and showed no defect in gingipain processing with a similar level of gingipain activity outside of the cell as for the Δ*porQ* and Δ*porZ* mutants. This suggests that PorF may also have a role during the latter stages of secretion, and as a predicted TonB-dependent receptor plug domain-containing protein (UNIPROT ID: Q7MWR1), our data supports a function in importing glycans involved in the A-LPS biogenesis and/or its linkage to secreted cargo (58).

**Table 3:** Summary of *P. gingivalis T9SS OMP* mutant phenotypes. ^†^Major (***), medium (**), or no (-) change compared with WT W50 phenotype. ^‡^Mutants with similar patterning are numbered the same. Unique patterning is numbered zero.

| *por*<br>KO | Colony<br>pigment<sup>†</sup> | Gingipain activity<sup>†</sup> |  |  |  | Immunoblots |  |  |  |  |  | OMV<br>size | OMV<br>size<br>range | Clust |
| --- | --- | --- | --- | --- | --- | --- | --- | --- | --- | --- | --- | --- | --- | --- |
|  |  | whole cell |  | supernatant |  | whole cell |  |  | supernatant |  |  |  |  |  |
|  |  | Rgp-A/B | Kgp | Rgp-A/B | Kgp | Rgp-A/B<br>‡ | Rgp-B<br>CTD<sup>‡</sup> | A-LPS | Rrg-A/B<br>‡ | Rrg-B<br>CTD<sup>‡</sup> | A-LPS |  |  |  |
| V | *** | *** | *** | *** | *** | 1 | 1 | ** | 0 | 0 | ** | *** | *** | 1 |
| U | *** | - | *** | - | *** | 0 | 0 | ** | 0 | 0 | ** | *** | *** | 2 |
| Q | ** | ** | ** | ** | ** | 2 | 2 | *** | 2 | 2 | *** | *** | *** | 3 |
| Z | ** | ** | ** | ** | ** | 2 | 2 | *** | 2 | 2 | *** | *** | *** | 3 |
| P | *** | *** | *** | *** | *** | 1 | 1 | ** | 1 | 1 | ** | *** | *** | 4 |
| T | *** | *** | *** | *** | *** | 1 | 1 | ** | 1 | 1 | ** | *** | *** | 4 |
| G | *** | *** | *** | *** | *** | 1 | 1 | ** | 1 | 1 | ** | *** | *** | 4 |
| F | ** | ** | ** | ** | ** | 0 | 0 | ** | 0 | 0 | ** | *** | *** | 5 |

We previously demonstrated that a W50 Δ*porV* mutant was unable to produce non-phosphorylated lipid A, (42) and we have now shown that this is also the case for all T9SS OMP mutants in W50. This implies that a functioning T9SS, rather than PorV alone, is required for LpxE lipid A 1-phosphatase activity. Likewise, all mutants produced larger deformed OMVs, comparable to those from a Δ*lpxE* mutant strain, when analysed by negative stain TEM. This has been previously observed for Δ*porU* and Δ*porT* mutants in W50 (42, 54) but also a Δ*rgpB*, but not Δ*rgpA* or Δ*kgp* mutant (42). Here we measured some reduction in RgpB activity, but not Kgp, in W50 Δ*lpxE*, and these observations could indicate a role for RgpB in directly or indirectly regulating LpxE. However, using NanoSight, we also measured two distinct forms of OMV in W50, W83 and 33277: a smaller well-defined species and a larger species with a broader range of diameters. In all our mutants we saw that the smaller species disappeared while the larger species remained. Single particle tracking has also been used to monitor OMV size distribution in the *P. gingivalis* strain 381, and has shown that WT 381 again produces both a smaller species and a broader range of vesicle sizes with diameters from ∼100 nm, and ∼150 to 400 nm, respectively (60). However, in a Δ*ppaD* mutant strain deficient in peptidylarginine deiminase (PPAD) activity, there was complete loss of the larger OMV species. PPAD is another cargo of the T9SS, and it appears that the T9SS may regulate the formation of both smaller and larger OMVs. It may also indicate that *P. gingivalis* the smaller species is dependent on regulated lipid-A modification, while the larger species form through an alternative pathway.

Many bacteria express lipid A modifying enzymes, but *P. gingivalis* is unusual as it produces both a lipid A 4-phosphatase (LpxE), two lipid A 4-phosphatases (LpxF1, LpxF2) and a 3-*O*-deacylase (LpxR). Furthermore, a triacylated lipid A species has also been observed in W50 OMVs, which is not detected in the Δ*porV* mutant, although no PagL-like 3-*O*-deacylase gene is present in *P. gingivalis* and this likely represents a novel enzyme (42). Our Δ*lpxE* mutant displayed a different protein profile in the outer membrane and OMVs compared with WT W50 and suggests in addition to LpxE being important for OMV blebbing, it also has a role in cargo sorting into OMVs. Moreover, we have identified a *P. gingivalis* like LpxE sequence with an extended C-terminal region in 21 additional species (**Supplementary Table 3**). Although sequence conservation is low in this region (**Supplementary Figure 1**), of those identified, *T. forsythia* and the Parabacteroides (*P. faecis*, *P. goldsteinii*, *P. timonensis*) and Porphyromonas (*P. gulae*, *P, loveana*) species also contain a *porV* gene and produce OMVs. This indicated that lipid-A modification may be a more general mechanism in other *Bacteroides* to couple type IX cargo sorting with OMV blebbing.

## Author contribution

Conceived and designed the experiments: S.W., Y.L., B.Z., J.A-O., R.B., G.M., P.M., M.R., M.C., J.G. Performed the experiments: S.W., Y.L., B.Z., J.A-O., R.B., G.M., P.M. Analyzed the data: S.W., Y.L., B.Z., J.A-O., R.B., G.M., P.M., M.R., M.C., J.G. Contributed reagents/materials/analysis tools: M.C., J.G. Wrote the paper: S.W., Y.L., B.Z., J.A-O., R.B., G.M., P.M., M.R., M.C., J.G.

## Acknowledgements

S.W., Y.L. and B.Z. were supported by China Scholarship Council.

## Supplementary Data

**Supplementary Figure 1:**
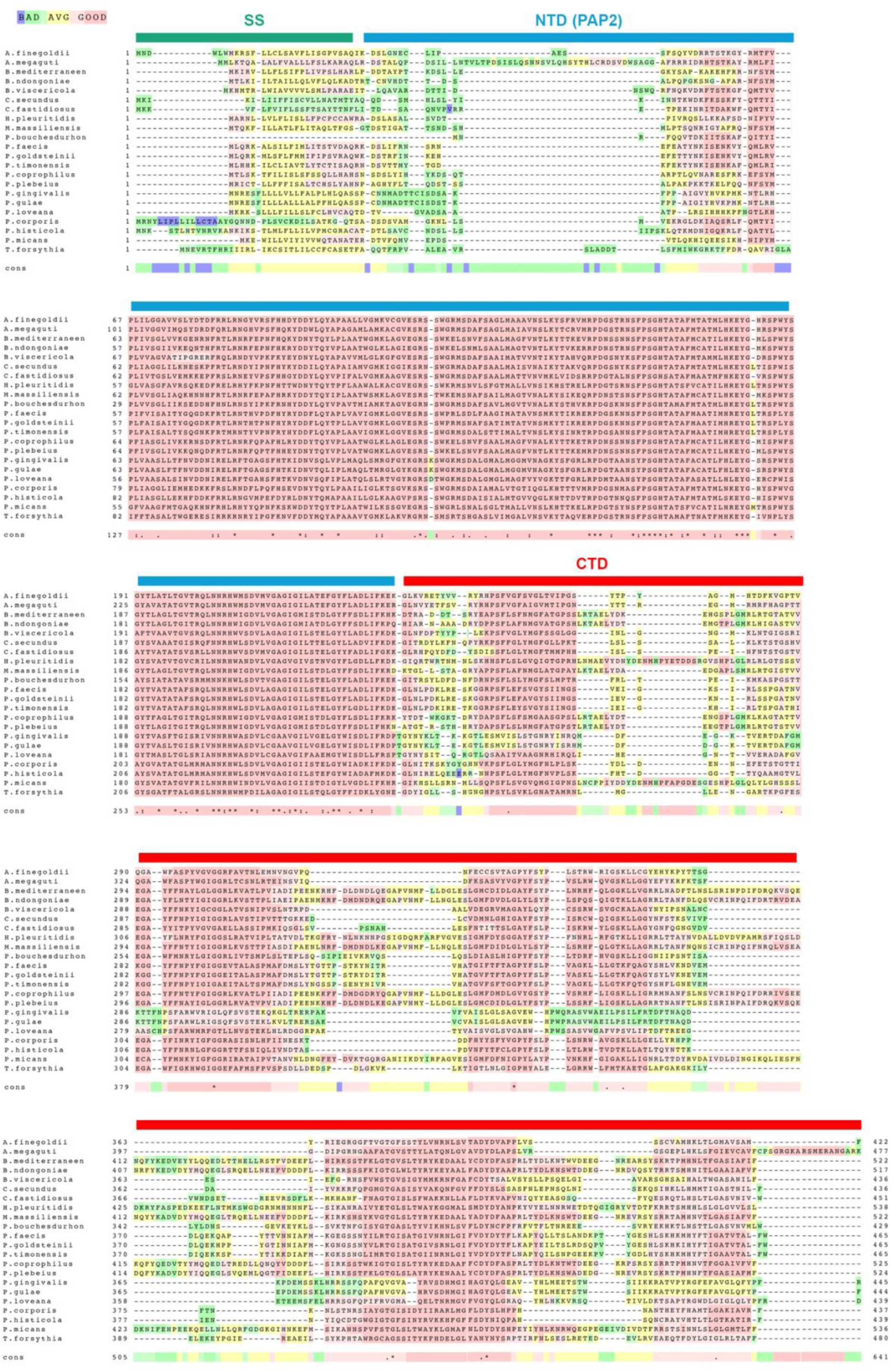
Multiple sequence alignment of LpxE homologs with a C-terminal extension. Reliability of the M-coffee (1) alignment is shown as blue, green, yellow, pink (bad to good). Fully conserved (*) are highlighted, as are residues with strongly (:) and weakly (.) similar properties.

**Supplementary Table 1:** Strains used in this study.

| Bacterial strain | Serogroup/ genotype | Reference |
| --- | --- | --- |
| *P. gingivalis* W50 | Wild type, no resistance. | (2) |
| *P. gingivalis* ΔporV | W50 porV (PG0027, PGN_0023) knock-out mutant, Clir. | (3) |
| *P. gingivalis* ΔporU | W50 porU (PG0026, PGN_0022) knock-out mutant, Clir. | This study |
| *P. gingivalis* ΔporQ | W50 porQ (PG0602, PGN_0645) knock-out mutant, Clir. | This study |
| *P. gingivalis* ΔporZ | W50 porZ (PG1604, PGN_0509) knock-out mutant, Clir. | This study |
| *E. coli* BL21 (DE3) | fluA2 [lon] ompT gal ($\lambda$ DE3) [dcm] ΔhsdS $\lambda$ DE3 = $\lambda$ sBamHIo ΔEcoRI-B int::(*lacI*::PlacUV5::T7 gene1) i21 Δin5 | NEB |
| *P. gingivalis* ΔporZ | W50 porZ (PG1604, PGN_0509) knock-out mutant, Clir. | This study |
| *P. gingivalis* ΔporP | W50 porP (PG0287, PGN_1677) knock-out mutant, Clir. | This study |
| *P. gingivalis* ΔporT | W50 porT (PG0751, PGN_0778) knock-out mutant, Clir. | This study |
| *P. gingivalis* ΔporG | W50 porG (PG0189, PGN_0297) knock-out mutant, Clir. | This study |
| *P. gingivalis* ΔporF | W50 porF (PG0534, PGN_1437) knock-out mutant, Clir. | This study |
| *P. gingivalis* Δkpg | W50 kpg (PGN_1728) knock-out mutant, Clir. | (4) |
| *P. gingivalis* ΔlpxE | W50 lpxE (PG1773, PGN_1713) knock-out mutant, Clir. | This study |
| *E. coli* BL21 (DE3) | Competent cells for protein expression, routine T7 expression. | NEB |

**Supplementary Table 2:**
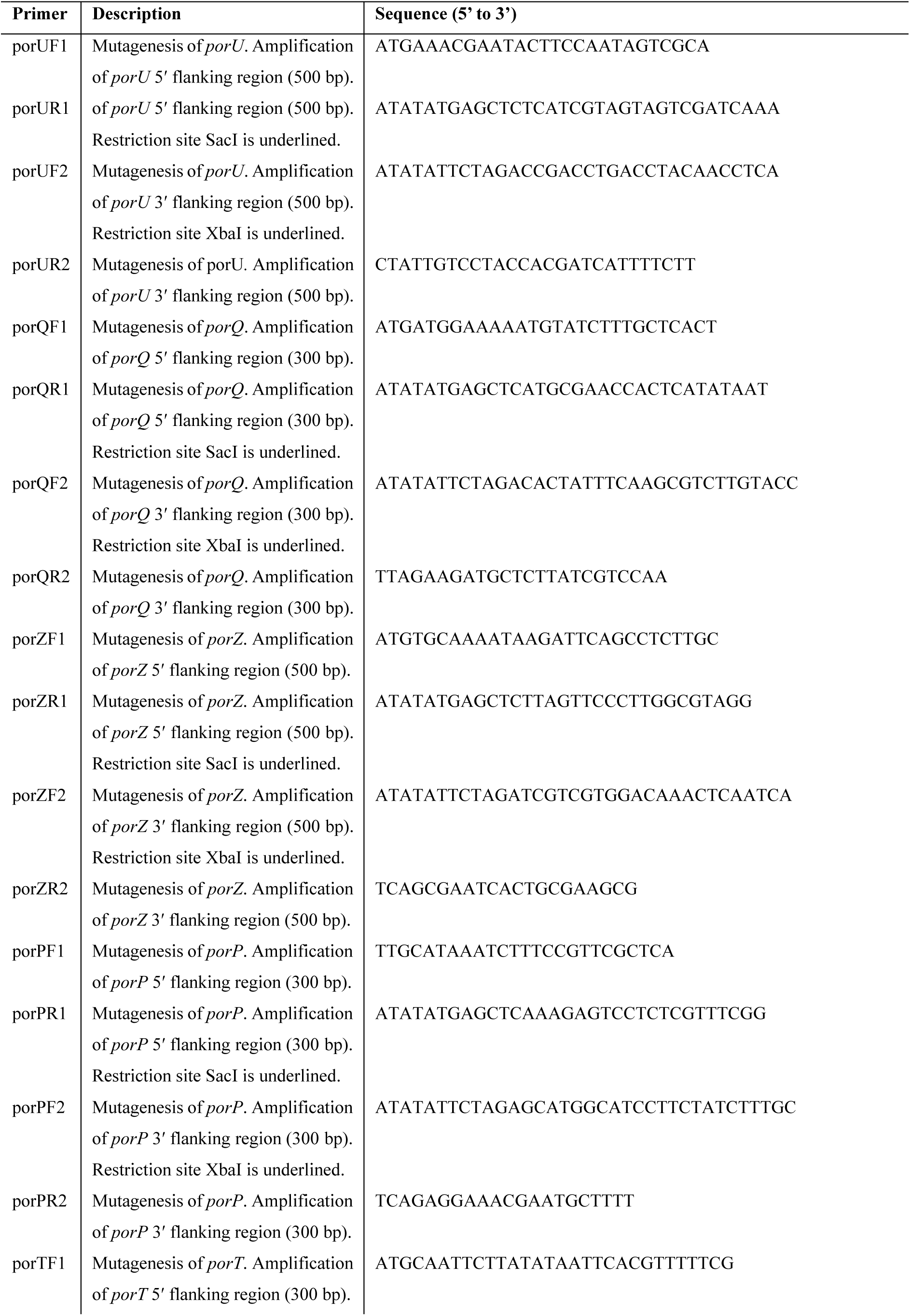
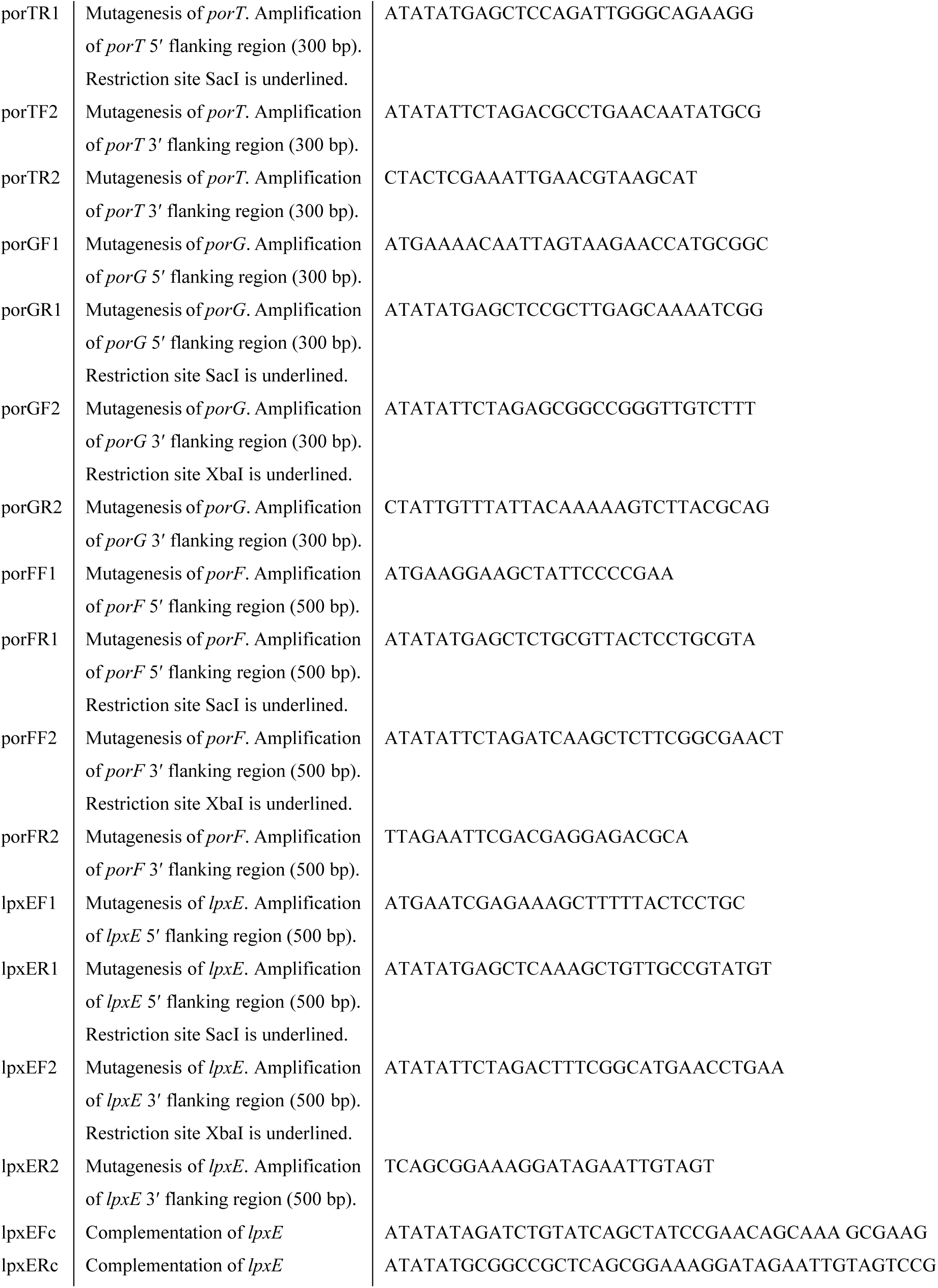
Primer used in this study.

**Supplementary Table 3:** LpxE homologs with extended C-terminal regions.

| **Description** | **Species** | **NIH accession code** |
| --- | --- | --- |
| phosphatase PAP2 family protein | *Alistipes finegoldii* | MBD9128102.1 |
| phosphatase PAP2 family protein | *Alistipes megaguti* | WP_232009156.1 |
| phosphatase PAP2 family protein | *Bacteroides mediterraneensis* | WP_083581745.1 |
| phosphatase PAP2 family protein | *Bacteroides ndongoniae* | WP_277122718.1 |
| phosphatase PAP2 family protein | *Barnesiella viscericola* | WP_289539706.1 |
| phosphatase PAP2 family protein | *Coprobacter secundus* | WP_021931000.1 |
| phosphatase PAP2 family protein | *Coprobacter fastidiosus* | WP_302545045.1 |
| phosphatase PAP2 family protein | *Hoylesella pleuritidis* | WP_021583963.1 |
| phosphatase PAP2 family protein | *Mediterranea massiliensis* | WP_289590024.1 |
| phosphatase PAP2 family protein | *Parabacteroides bouchesdurhonensis* | WP_102409487.1 |
| phosphatase PAP2 family protein | *Parabacteroides faecis* | WP_258915275.1 |
| phosphatase PAP2 family protein | *Parabacteroides goldsteinii* | WP_258971860.1 |
| phosphatase PAP2 family protein | *Parabacteroides timonensis* | WP_075559510.1 |
| phosphatase PAP2 family protein | *Phocaeicola coprophilus* | WP_278710372.1 |
| phosphatase PAP2 family protein | *Phocaeicola plebeius* | WP_304122999.1 |
| lipid A C1-phosphatase | *Porphyromonas gingivalis* | WP_005873717.1 |
| lipid A C1-phosphatase | *Porphyromonas gulae* | WP_039437269.1 |
| phosphatase PAP2 family protein | *Porphyromonas loveana* | WP_116678422.1 |
| phosphatase PAP2 family protein | *Prevotella corporis* | WP_277265612.1 |
| phosphatase PAP2 family protein | *Prevotella histicola* | WP_278527811.1 |
| phosphatase PAP2 family protein | *Prevotella micans* | WP_006953444.1 |
| phosphatase PAP2 family protein | *Tannerella forsythia* | WP_080948576.1 |

## Notes

### Competing Interest Statement

The authors have declared no competing interest.

